# Promotion of hyperthermic-induced rDNA hypercondensation in *Saccharomyces cerevisiae*

**DOI:** 10.1101/838821

**Authors:** Donglai Shen, Robert V. Skibbens

## Abstract

Ribosome biogenesis is tightly regulated through stress-sensing pathways that impact genome stability, aging and senescence. In *Saccharomyces cerevisiae*, ribosomal RNAs are transcribed from rDNA located on the right arm of chromosome XII. Numerous studies reveal that rDNA decondenses into a puff-like structure during interphase and condenses into a tight loop-like structure during mitosis. Intriguingly, a novel and additional mechanism of increased mitotic rDNA compaction (termed hypercondensation) was recently discovered that occurs in response to temperature stress (hyperthermic-induced) and is rapidly reversible. Here, we report that neither changes in condensin nor cohesin binding dynamics appear to play a critical role in hyperthermic-induced rDNA hypercondensation – differentiating this architectural state from normal mitotic condensation (requiring cohesins and condensins) and the premature condensation (requiring condensins) that occurs during interphase in response to nutrient starvation. A candidate genetic approach revealed that deletion of either Hsp82 or Hsc82 (Hsp90 heat shock paralogs) result in significantly reduced hyperthermic-induced rDNA hypercondensation. Intriguingly, Hsp inhibitors do not impact rDNA hypercondensation. In combination, these findings suggest that Hsp90 either stabilizes client proteins, which are sensitive to very transient thermic challenges, or directly promotes rDNA hypercondensation during preanaphase. Our findings further reveal that the high mobility group protein Hmo1 is a negative regulator of mitotic rDNA condensation, distinct from its role in promoting premature-condensation of rDNA during interphase upon nutrient starvation.

## INTRODUCTION

Protein synthesis in all organisms takes place in the highly-conserved ribonucleoprotein complex - the ribosome. Ribosome biogenesis is thus directly related to cell growth and proliferation (Kief and Warner, 1981). In eukaryotes, the nuclear compartment that assembles ribosomes (including rRNA synthesis, processing and ribonucleoparticle assembly), is termed the nucleolus. rRNA arises from transcription of the rDNA locus that resides on the right arm of chromosome XII in the *Saccharomyces cerevisiae* yeast genome. This locus is approximately 1-2 Mb and consists of about 150 tandem repeats, each of which is 9.1 kb and encodes for 5S, 5.8S, 25S, and 18S rRNAs (Spahn et al., 2001; Nomura, 2001; Sirri et al., 2008).

Alterations in rDNA structure and function have implications far beyond the canonical roles of the nucleolus in rDNA transcription and ribosome biogenesis (Tiku and Antebi, 2018; Schöfer and Weipoltshammer, 2018; Kobayashi and Sasaki, 2017). For instance, rDNA is the most highly represented gene in any eukaryote and also the most heavily transcribed locus (accounting for over 60% of the entire RNA pool) (Tomson et al., 2006). Due to this highly repetitive structure and active transcriptional status, rDNA is the most recombinogenic, and therefore mutagenic, site within the eukaryotic genome (Nomura, 2001; Tomson et al., 2006; Pal et al., 2018; Kwan et al., 2016). The importance of maintaining rDNA locus stability is highlighted by the fact that DNA replication forks are programmed to stall within rDNA, precluding catastrophic head-on collision of replication and transcription complexes (Biswas et al., 2017; Weitao et al., 2003; Shyian et al., 2016). Furthermore, rDNA transcription rates, and even nucleolar size, are intimately coupled to changes in nutrient levels, revealing that rDNA plays a central role in responding to environmental cues (Li et al., 2006; Tsang et al., 2007; Wang et al., 2016). Disruption of rDNA transcription leads to ribosome biogenesis stress and also inhibits Mdm2 function, resulting in cell cycle arrest, senescence, and apoptosis through p53-dependent pathways (Turi et al., 2018).

In yeast, changes in rDNA homeostasis impacts cellular aging and replicative lifespan in which extrachromosomal rDNA circles (ERCs), that arise through recombination, deplete the remaining genome of critical regulatory factors (Lewinska et al., 2014; Park et al., 1999; Sinclair and Guarente, 1997; Sinclair et al., 1997; Shcheprova et al., 2008). Clinically, disruption of rDNA function in humans results in neurodegeneration, tumorigenesis and severe developmental defects that include Treacher-Collins Syndrome, Blackfan Anemia, CHARGE Syndrome and several others (Danilova and Gazda, 2015; Udugama et al., 2018; Wang and Lemos, 2017; Xu et al., 2014; Xu et al., 2017; Hallgren et al., 2014). Given that a rather surprisingly small percentage of nucleolar proteins function in ribosome biogenesis (Tiku and Antebi, 2018; Schöfer and Weipoltshammer, 2018), it becomes critical to explore the regulatory mechanisms through which rDNA responds to the many challenges imposed on the cell to ensure proper development and cell cycle regulation.

rDNA structure is tightly regulated through the cell cycle. In budding yeast, rDNA forms a diffuse puff-like structure during G1 phase that coalesces into a tight loop-like structure during mitosis (Guacci et al., 1994; Guacci et al., 1997). The importance of these architectural changes is highlighted by the development of numerous strategies that include FISH, GFP-tagged rDNA binding proteins, and a streamlined intercalating-dye method which now provides for rapid and efficient quantification of rDNA condensation (Guacci et al., 1994; Guacci et al., 1997; D’Ambrosio et al., 2008; Lopez-Serra et al., 2013; Tong and Skibbens, 2015; Lavoie et al., 2002; Lavoie et al., 2004; Shen and Skibbens, 2017a). To date, these condensation assays have helped elucidate the role of highly conserved cohesin and condensin complexes in regulating rDNA architecture. This is due to the fact that mutations in every cohesin and condensin subunit tested, or mutation of cohesion regulators such as the cohesin loader Scc2-Scc4 and cohesin acetyltransferase Eco1/Ctf7 (herein Eco1), produce profound impacts on condensation such that rDNA fails to compact during mitosis and appears instead as diffuse puff-like structures (D’Ambrosio et al., 2008; Lopez-Serra et al., 2013; Tong and Skibbens, 2015; Skibbens et al., 1999; Tóth et al., 1999; Kueng et al., 2006; Hirano, 2012). In addition to appropriate condensation reactions that occur during mitosis, the rDNA locus can also condense during G1 phase in response to nutrient starvation or rapamycin treatment. This premature rDNA condensation, which includes nucleolar contraction, requires *de novo* recruitment of condensin and the high mobility group protein Hmo1 (Tsang et al., 2007; Wang et al., 2016).

Despite the intense focus on yeast rDNA architecture over the last two decades (Xu et al., 2017; Guacci et al., 1994; Lavoie et al., 2004; Freeman et al., 2000; Castaño et al., 1996; Sullivan et al., 2004; Gard et al., 2009; Hong and Seong, 2014), an additional rDNA state was only recently discovered in which mitotic cells induce a hyper-condensed rDNA state in response to elevated temperature (Shen and Skibbens, 2017a; Matos-Perdomo and Machín, 2018a). This hyperthermic-induced rDNA hypercondensation is both rapidly induced and reversible. Intriguingly, rDNA hypercondensation is also inducible by numerous cell stressors (rapamycin exposure, oxidative stress, nitrogen starvation, and caloric restriction) that inhibit the TORC1 pathway (Matos-Perdomo and Machín, 2018a,b). The extent to which hyperthermic-induced rDNA hypercondensation is predicated on cohesin or condensin dynamics, however, remains unknown. Here, we find that, unlike the changes in either cohesin or condensin dynamics required for mitotic condensation, hyperthermic-induced rDNA hypercondensation occurs in the absence of altered levels of either condensin or cohesin. Instead, we find that mutation of heat shock/chaperone Hsp90 family members Hsp82 and Hsc82 result in significantly reduced rDNA hypercondensation. Our results further identify Hmo1 as a negative regulator of mitotic rDNA condensation, in opposition to its role in rDNA premature-condensation that occurs during interphase upon nutrient starvation (Tsang et al., 2007; Wang et al., 2016).

## RESULTS

### Hyperthermic-induced rDNA hypercondensation occurs in the absence of new cohesin deposition (Scc2 inactivation) and release (Rad61 deletion)

Wildtype cells shifted to an elevated temperature during mitosis exhibit rDNA hypercondensation (Shen and Skibbens, 2017a; Matos-Perdomo and Machín, 2018a), but the structural basis for this dramatic change in chromatin structure remains unknown. Cohesins play a critical role in chromosome condensation, including across the rDNA locus, such that mutation in either cohesin subunits (Mcd1/Scc1, Pds5 or Scc3) or regulators (Eco1 or Scc2) all result in severe rDNA condensation defects (Guacci et al., 1997; D’Ambrosio et al., 2008; Tong and Skibbens, 2015; Skibbens et al., 1999; Hartman et al., 2000; Woodman et al., 2015; Orgil et al., 2015). These observations formally suggest that *de novo* cohesin deposition during mitosis may play a critical role in hyperthermic-induced rDNA hypercondensation, in contrast to the decondensation of rDNA into ‘puffs’ that occurs upon either cohesin inactivation or dissociation (Guacci et al., 1997; Ciosk et al., 2000; Shen and Skibbens, 2017a). Here, we test whether *de novo* cohesin deposition promotes hyperthermic-induced rDNA hypercondensation by inactivating the Scc2,4 heterocomplex that is required for cohesin deposition onto DNA (Ciosk et al., 2000; Watrin et al., 2006). Log phase cultures of wildtype and *scc2-4* mutant cells were synchronized in G1 at 23°C using rich medium supplemented with alpha factor, washed and then arrested in preanaphase at 23°C (permissive for *scc2-4* cells) by incubation in medium supplemented with nocodazole. The resulting cultures were then shifted to 37°C (non-permissive for *scc2-4* cells) for 1 hour while maintaining the preanaphase arrest. Cell cycle progression from log phase into mitosis was confirmed by flow cytometry (Figure 1A). As expected, mitotic wildtype cells maintained at 23°C contained long rDNA loops while the rDNA of mitotic cells shifted to 37°C during the final hour of incubation hypercondensed into very short loops (Figure 1B), consistent with prior findings (Shen and Skibbens, 2017a; Matos-Perdomo and Machín, 2018a). Prior analyses of these cells revealed that only a fraction of *scc2-4* mutant cells contain condensed rDNA loci at 23°C, a level which is retained after shifting to 37°C during the final hour of incubation (Shen and Skibbens, 2017b). We thus limited our current measurements to the fraction of cells in which rDNA loops were tightly cohered and condensed into discrete loops (Figure 1B). The results show that *scc2-4* cells contain long rDNA loops at 23°C. Importantly, rDNA in *scc2-4* mutant cells fully hypercondense into very short loops after incubation at 37°C for 1 hour (Figure 1B, C). We confirmed that this *scc2-4* mutant strain is indeed temperature sensitive and defective in cohesin deposition onto chromatin (Shen and Skibbens, 2017b). Thus, temperature-induced rDNA hypercondensation during mitosis occurs in the absence of *de novo* cohesin deposition.

**Figure 1.**
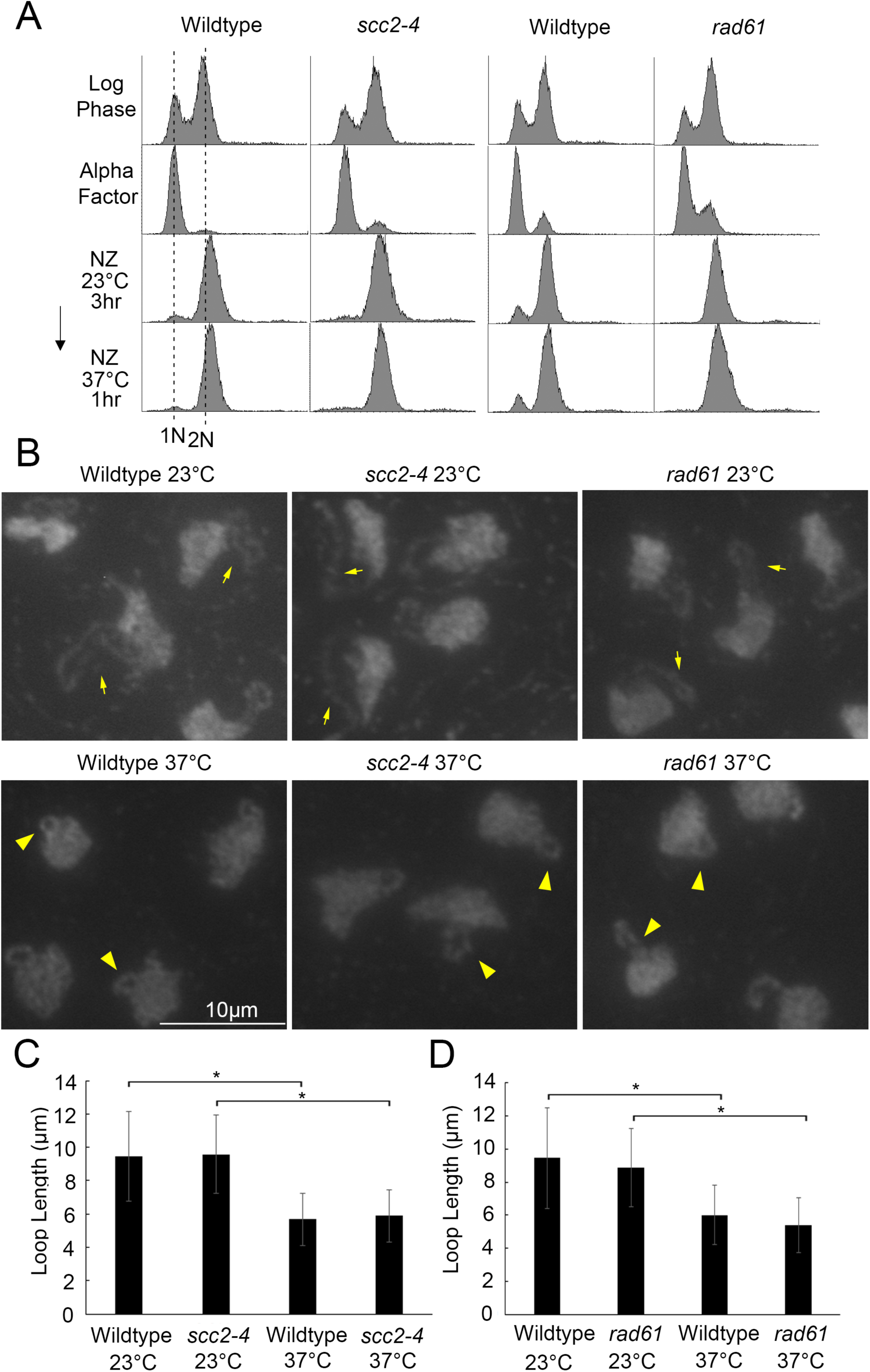
Cohesin deposition and/or release are not required for hyperthermic-induced rDNA hypercondensation. A) Flow cytometer data documents DNA content of Wildtype (YBS1039), *scc2-4* mutant cells (YMM511) and *rad61* null cells (YBS2037) throughout the experiment. Wildtype and *scc2-4* mutant cells were processed separately from wildtype and *rad61* null cells. Cells were maintained in nocodazole for 3 hours at 23°C, post-alpha factor arrest, followed by an additional 1 hour incubation at 37°C. B) Chromosomal mass and rDNA loop structures detected using DAPI. Yellow arrows indicate long rDNA loops. Yellow arrowheads indicate short rDNA loops. All field of views are shown at equal magnification. C) Quantification of the loop length of condensed rDNA in wildtype and *scc2-4* mutant cells. Data shown was obtained from 3 biological replicates. For Wildtype versus *scc2-4* mutant, the results reflect n values of 228 of Wildtype cells at 23°C, 235 for *scc2-4* cells at 23°C, 214 for Wildtype cells at 37°C, and 201 for *scc2-4* cells at 37°C. D) Quantification of loop lengths of condensed rDNA in wildtype and *rad61* null cells. For Wildtype versus *rad61* null mutant, the results reflect n values of 235 for Wildtype cells at 23°C, 214 for *rad61* cells at 23°C, 157 for Wildtype cells at 37°C, and 151 for *rad61* cells at 37°C. Statistical analysis was performed using a Tukey HSD one way ANOVA. P-Value = 0.721 indicates that there is no significant difference between the average loop lengths of wildtype cells (5.67 μm) and *scc2-4* mutant cells (5.89 μm) at 37°C in panel C. P-Value = 0.103 indicates there is no significant difference between wildtype cells (6.02 μm) and *rad61* null cells (5.40 μm) at 37°C in panel D. Statistical significant differences (*) are based on P < 0.05.

Wildtype cells exhibit faster growth kinetics at 37°C, despite containing hypercondensed rDNA (Shen and Skibbens, 2017a; Matos-Perdomo and Machín, 2018a). To accommodate the increase in rDNA transcription required for this faster rate of cell growth, we previously posited that rDNA hypercondensation (axial loop shortening or decreased loop contour length) might occur through cohesin dissociation to promote increased lateral looping (Shen and Skibbens, 2017a). To test whether cohesin removal promotes shorter axial rDNA loops (interpreted as hypercondensation), we turned to the cohesin destabilizer Rad61/WAPL (Kueng et al., 2006; Lopez-Serra et al., 2013; Game et al., 2003; Vernì et al., 2000; Sutani et al., 2009). Log phase wildtype and *rad61* null cells were treated as described earlier to achieve sequential G1 and preanaphase synchronizations at 23°C before shifting to 37°C for 1 hour, while maintaining the mitotic arrest (Figure 1A). *rad61* null cells condensed the rDNA into extended discrete loops at 23°C, similar to both wildtype and *scc2-4* mutant cells. Moreover, *rad61* null cells were fully competent to hypercondense the rDNA into very short loops upon shifting to 37°C for 1 hour (Figure 1B, D). These results reveal that rDNA hypercondensation (axial shortening) occurs in the absence of Rad61-dependent cohesin release from DNA. In combination, these results reveal that mitotic hyperthermic-induced rDNA hypercondensation occurs independent of both Scc2-dependent deposition of new cohesins and Rad61-dependent release of pre-existing cohesins.

Chl1 DNA helicase is a positive regulator of both sister chromatid cohesion and chromosome condensation in that Chl1 promotes both Scc2 and cohesin binding to DNA (Skibbens, 2004; Mayer et al., 2004; Inoue et al., 2007; Xu et al., 2007; Laha et al., 2011; Rudra and Skibbens, 2012; Borges et al., 2013; Samora et al., 2016; Shen and Skibbens, 2017b), providing a further opportunity to assess whether cohesin deposition/release are involved in hyperthermic-induced rDNA hypercondensation. We previously reported analyses of rDNA loop lengths in wildtype cells (adapted from Figure 1E in Shen and Skibbens, 2017a), but at that time did not include quantification of rDNA loop lengths in *chl1* mutant cells that were simultaneously assessed. As previously described, wildtype and *chl1* deletion cells were synchronized in G1 at 23°C, then cultures divided and released into either 23°C or 37°C medium supplemented with nocodazole to arrest cells in preanaphase. Cell cycle progression and arrests were confirmed using flow cytometry (Supp. Figure 1A). Our results reveal that *chl1* null cells, shifted to 37°C, were fully competent to hypercondense the rDNA into very short loops, similar to wildtype cells and in contrast to the elongated rDNA loops present at 23°C in both wildtype and *chl1* mutant cells (Supp Figure 1B, C and Shen and Skibbens, 2017a). Thus, hyperthermic-induced rDNA hypercondensation occurs in the absence of Chl1, consistent with a mechanism independent of both Scc2 and cohesin dynamics.

### Condensin deposition and/or release are not required for hyperthermic-induced rDNA hypercondensation

Mitotic chromosome condensation requires condensin, in addition to cohesin, such that condensin mutants exhibit severe condensation defects along the rDNA locus (D’Ambrosio et al., 2008; Lavoie et al., 2002; Lavoie et al., 2004; Freeman et al., 2000; Strunnikov et al., 1995). Unlike the cohesin complex, there is no known loading complex that promotes condensin deposition onto chromosomes (Hirano, 2012; Ganji et al., 2018). Thus, to assess whether condensin deposition is required for hyperthermic-induced rDNA hypercondensation, we directly tested for hyperthermic-induced changes in condensin binding to rDNA using chromatin immunoprecipitation (ChIP). Wildtype cells expressing HA-tagged Smc2 were synchronized in G1 at 23°C, then divided into two with aliquots released into either 23°C or 37°C fresh medium supplemented with nocodazole to arrest cells in preanaphase (Figure 2A). Protein-DNA complexes were cross-linked using formaldehyde, followed by cell lysis and sonication to shear the DNA. Chromatin complexes containing Smc2 were immunoprecipitated, cross-links reversed and condensin enrichment quantified from PCR using four well-documented condensin-binding sites within the rDNA locus (Figure 2B) (Johzuka and Horiuchi, 2009; Thattikota et al., 2018). The results, averaged across all 4 sites and based on 3 independent biological replicates (at 23°C versus 37°C), reveal no change in Smc2 enrichment (p-value = 0.24), despite dramatic changes in rDNA structure. Even on a site-by-site analyses, the results suggest that Smc2 levels do not increase during rDNA hypercondensation but instead remain relatively unchanged at 23°C compared to 37°C (Figure 2C). Thus, hyperthermic-induced rDNA hypercondensation occurs independent of both condensin deposition and dissociation.

**Figure 2.**
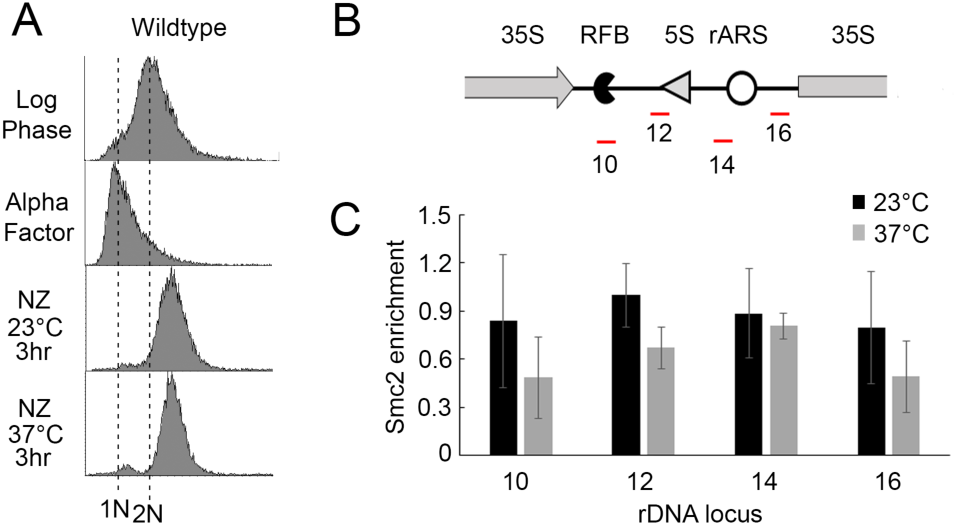
Condensin deposition and/or releasing appear independent of hyperthermic-induced rDNA hypercondensation. A) Flow cytometer data documents DNA content throughout the experiment. Wildtype cells (YBS3036) were maintained in nocodazole for 3 hours at 23°C or 37°C post-alpha factor arrest. B) Schematic indicates the location of four ChIP primer sets (in red) that reside within the interval region flanked by two rDNA repeats. RFB = Replication Fork Barrier; rARS = rDNA Autonomously Replicating Sequence. C) Quantification of ChIP using the primer pairs shown in (B) to assess Smc2 fold enrichment in mitotic wildtype cells maintained at 23°C versus 37°C resulting. Data shown was obtained from 3 biological replicates. Error bars represent standard deviation of each sample. Statistical analysis was performed using Tukey HSD one way ANOVA. P-Value = 0.694, 0.752, 0.899, 0.815 for primer sets 10, 12, 14, 16 respectively, and indicate that there is no significant difference between the Smc2-HA enrichment of wildtype cells arrested at 23°C and 37°C. Note that combining data obtained from the 4 sites, and then comparing the average Smc2 enrichment from 3 independent biological replicates at 23 degree to 37 degree using Student’s T test, produces a p-value of 0.24. This further indicates that, by testing for trends across the rDNA in aggregate, condensin deposition and/or release are not required for hyperthermic-induced rDNA hypercondensation. Statistical significant differences (*) are based on P < 0.05.

A recent study suggests that rDNA condensation and nucleolar compaction progress to a maximum state during early anaphase (de los Santos-Velazquez et al., 2017). While this rDNA condensation appears separate from the significant decrease in rDNA axial lengths that occur in response to heat-stress preanaphase (Shen and Skibbens, 2017a; Matos-Perdomo and Machín, 2018a), we decided to augment our arrest strategy to ensure that cells are not escaping the nocodazole-induced mitotic arrest. Cdc23 is an essential component of the Anaphase Promoting Complex (Hartwell et al., 1973; Lamb et al., 1994; Irniger et al., 1995). *cdc23-1* mutant strains were synchronized in G1 (alpha factor) and then released into 37°C (restrictive for *cdc23* alleles) medium supplemented with or without DMSO or nocodazole. DNA profiles confirm the efficacy of *cdc23* mutant protein inactivation in that 2N DNA profiles were obtained at the end of the 3 hour temperature incubation at 37°C regardless of the presence or absence of nocodazole or DMSO (Supp. Figure 2A). Nuclei in APC mutants cells, arrested preanaphase in the absence of nocodazole, experience mitotic forces via kinetochore microtubules and spindles. We have observed a large population of distort chromatin that result in discernible rDNA loop structures (Supp. Figure 2B, red arrows). Prior findings similarly reported that the strategies used to arrest cells preanaphase impact rDNA architectures (Guacci et al., 1994). More importantly, *cdc23-*1 cells exhibited highly hypercondensed rDNA under hyperthermic conditions (driving both rDNA axial shortening and APC inactivation) with or without nocodazole (Supp. Figure 2B, yellow arrows). These results confirm that rDNA hypercondensation can be induced prior to anaphase solely by increased temperature and is not a by-product of nocodazole treatment.

### Hyperthermic-induced rDNA hypercondensation is separate from several activities that impact rDNA regulation

The surprising finding that neither cohesin nor condensin dynamics contribute to hyperthermic-induced rDNA hypercondensation suggested that a novel mechanism must exist by which cells regulate rDNA structure in response to thermic stress. We thus turned to heat-shock pathways through which cells appropriately respond to elevated temperatures (Verghese et al., 2012), even though no evidence to date directly implicates heat shock proteins/chaperones (HSP/C) either in mitotic rDNA condensation or hyperthermic-induced hypercondensation. To generate a candidate list, we first took a bioinformatics approach and queried the Saccharomyces Genome Database (SGD) GO term database using an iterative process in which each search contained unique combinations of any two of several terms (Response to heat; Nucleolus; Chromatin binding; Regulation of DNA metabolic process, etc). We cross-referenced the resulting lists to identify candidates that occur in high frequency and then selected those in which mutations are readily obtainable from a prototrophic deletion collection (Winzeler et al., 1999; Giaever and Nislow, 2014; VanderSluis et al., 2014). We finally prioritized 10 genes that provide the most extensive coverage of independent heat shock/chaperone pathways (Table 1).

**Table 1.** Prioritized list of heat shock/chaperone encoding genes obtained from iterative GO terms searches and that represent a diverse set of cellular responses to elevated temperature.

| **Comm**<br>**on** | **Systematic** | **Descriptor** | **Referenc**<br>**e** |
| --- | --- | --- | --- |
| *FOB1* | YDR110W | rDNA<br>replication fork<br>barrier | SGD |
| *HIT1* | YJR055W | snoRNP<br>assembly factor | SGD |
| *HMO1* | YDR174W | High mobility<br>group factor | SGD |
| *HSP82* | YPL240C | Hsp90<br>chaperone | SGD |
| *ISW1* | YBR245C | Imitation-swi<br>tch chromatin<br>remodelers | SGD |
| *MSN2* | YMR037C | Stress-respo<br>nsive<br>transcriptional<br>activator | SGD |
| *MSN4* | YKL062W | Stress-respo<br>nsive<br>transcriptional<br>activator | SGD |
| *SIR2* | YDL042C | NAD+<br>dependent histone deacetylase | SGD |
| *SSA1* | YAL005C | ATPase<br>member of HSP70<br>family | SGD |
| *TOP1* | YOL006C | Topoisomer<br>ase I | SGD |

**Table 2.**
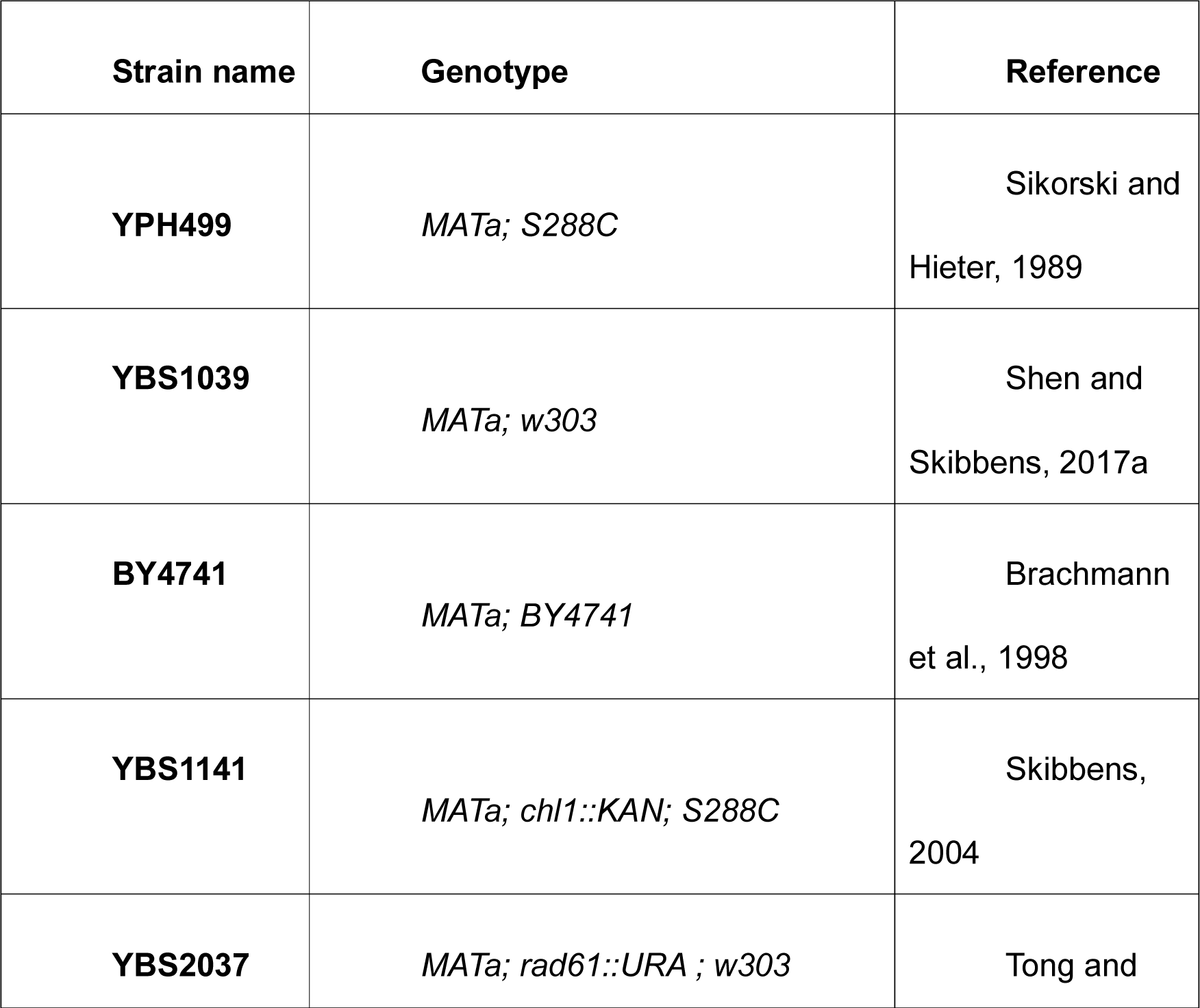
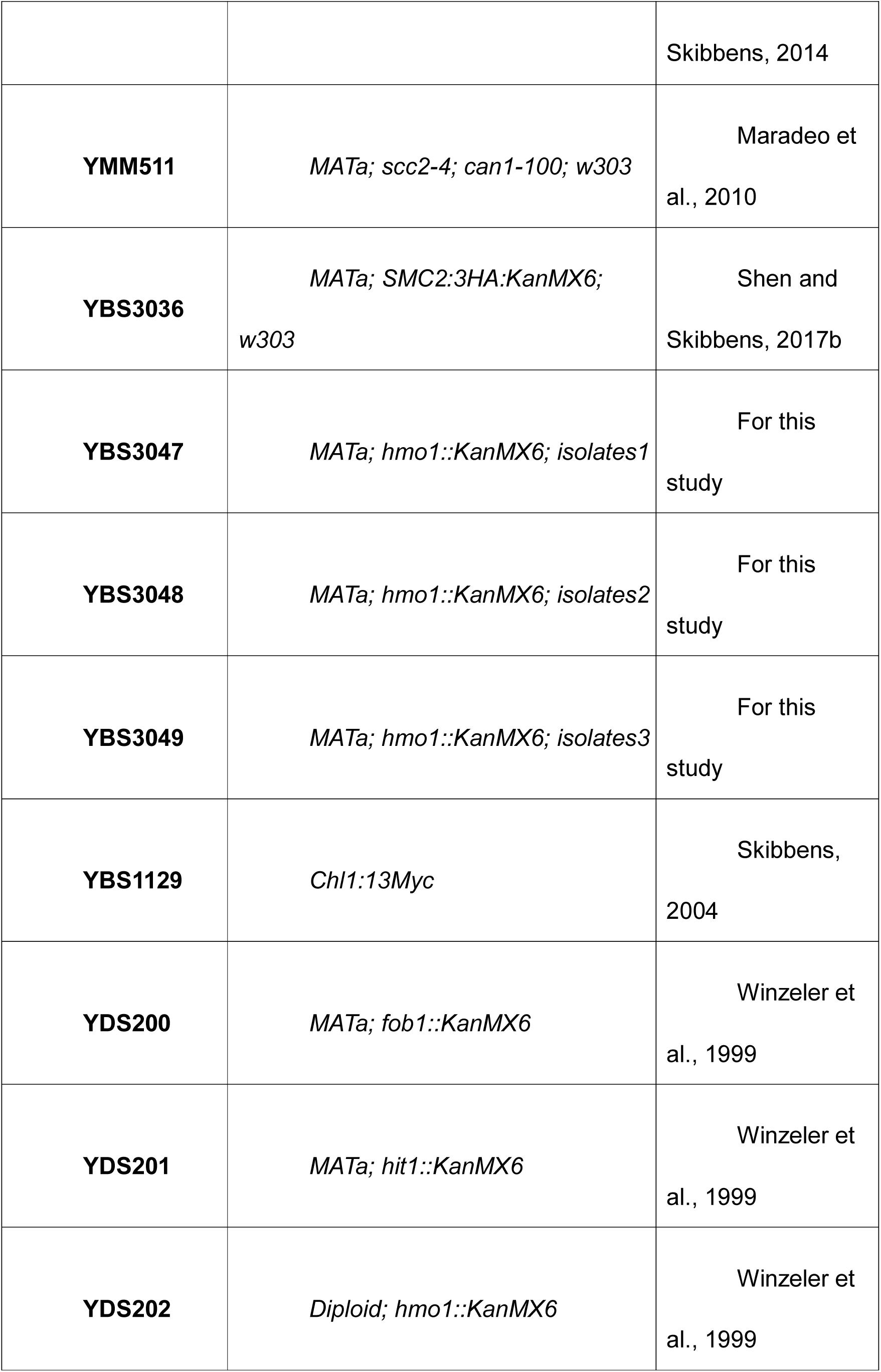
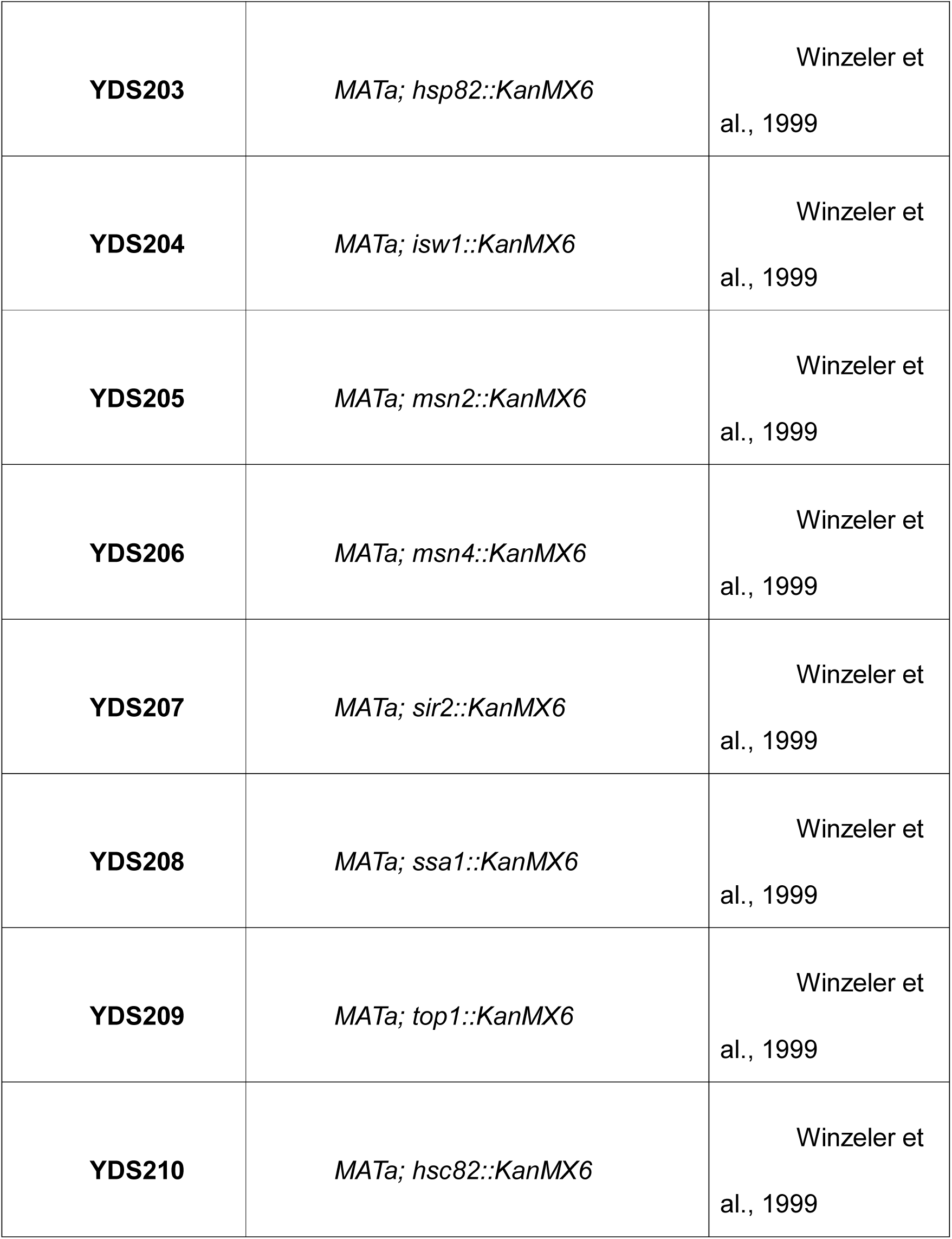
Yeast strains used in this study.

Wildtype and all 10 heat shock protein/chaperone (HSP/C) null cells were sequentially synchronized in G1 and preanaphase as described above, before shifting the resulting mitotic cells to 37°C for 1 additional hour while maintaining the mitotic arrest. Cell cycle synchronizations and progression for each strain was monitored using flow cytometry (Figure 3A). As expected, rDNA in wildtype cells exhibited significantly hypercondensed rDNA loops after shifting to 37°C for 1 hour (Figure 3B). Not surprisingly, the bulk of the HSP/C candidates (*msn2, msn4, ssa1, sir2, isw1, hit1* and *fob1*) exhibited both normal mitotic rDNA condensation at 23°C and hypercondensation at 37°C (Figure 3B). Thus, rDNA hyperthermic-induced hypercondensation is a specialized and unique response that is independent of most heat shock pathways. Of particular interest, the results reveal that neither deletion of the Sir2 (NAD+ deacetylase and major regulator of rDNA silencing and structure) nor Fob1 (the rDNA replication fork barrier protein that coordinates rDNA replication with transcription) had any adverse impact on rDNA hyperthermic-induced hypercondensation (Figure 3B) (Kobayashi and Sasaki, 2017; Johzuka and Horiuchi, 2009; Gottlieb and Esposito, 1989; Smith and Boeke, 1997; Fritze et al., 1997; Imai et al., 2000; Machín et al., 2004). These results highlight the physiological separation of these pathways from rDNA hypercondensation. Notably, *top1* null cells exhibited 44% of puff-like rDNA structures even at 23°C (similar condensation defects were observed at 37°C), indicating that the rDNA was decondensed regardless of temperature. Thus, *top1* was excluded from further analyses into the mechanism of hyperthermic-induced hypercondensation. Our results, however, document that Top1 is critical for rDNA condensation at all temperatures (Figure 3B), consistent with prior findings that *top1* promotes condensation in *Drosophila melanogaster* and that *top1* null cells exhibit rDNA condensation defects in budding yeast (Castaño et al., 1996; Zhang et al., 2000).

**Figure 3.**
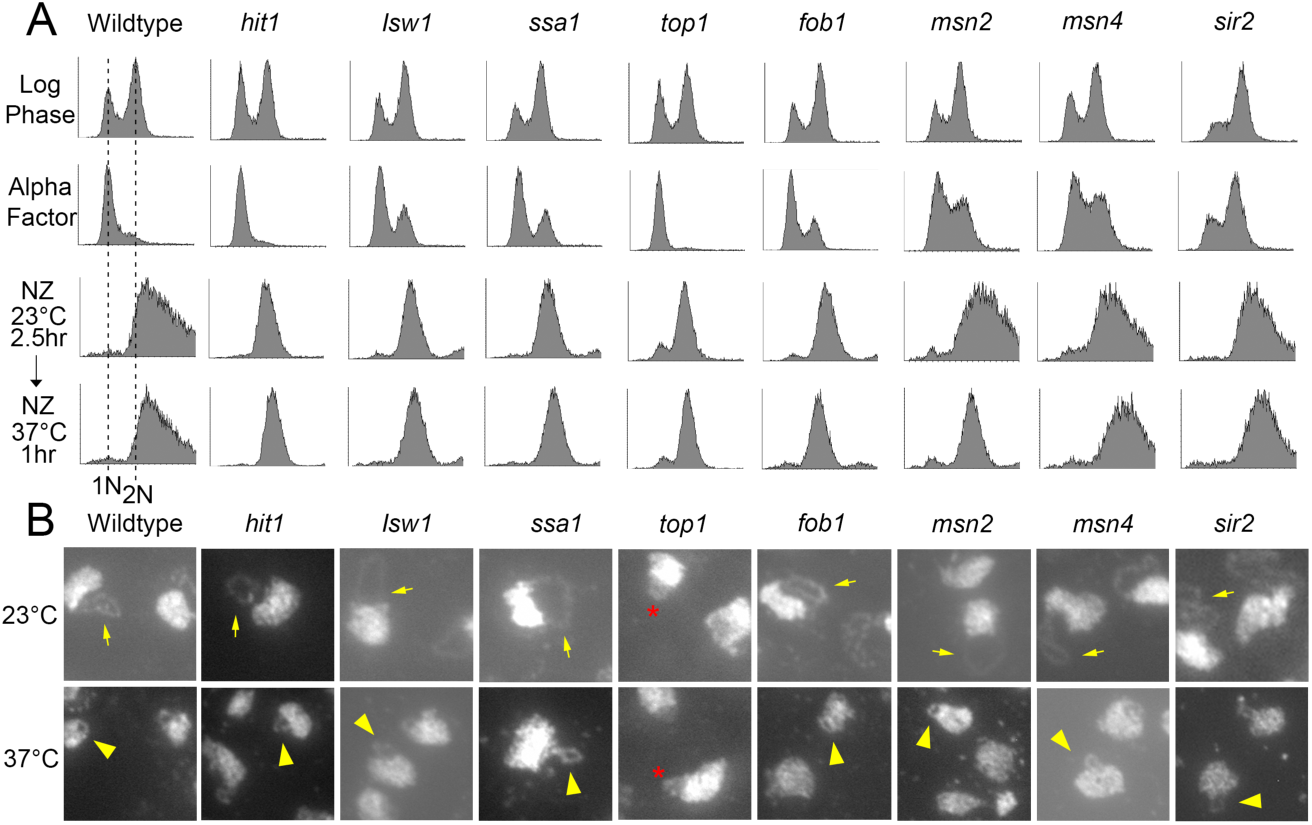
Hyperthermic-induced rDNA hypercondensation is separate from several activities that impact rDNA regulation. A) Flow cytometer data documents DNA contents of log phase cells (wildtype and indicated HSP/C mutant strains) synchronized in G1 (Alpha Factor), then subsequently arrested in preanaphase (NZ) at 23°C for 2.5 hours before shifting to 37°C for 1 hour. B) Chromosomal mass and rDNA loop structures detected using DAPI. Yellow arrows indicate long rDNA loops. Yellow arrowheads indicate short rDNA loops. Red star indicates the decondensed rDNA puff observed in *top1* null mutant.

### Hsp90 mutants are defective in hyperthermic-induced rDNA hypercondensation

Given the basis of our bioinformatics-based strategy, we were surprised to find an HSP/C that indeed impacts hyperthermic-induced rDNA condensation. Condensation assays in which hypercondensation was induced for a single hour during a mitotic arrest revealed that *hsp82* null cells fully support normal rDNA condensation during mitosis at 23°C, but failed to completely hypercondense rDNA to wildtype levels in response to 37°C incubation (Figure 4A, B). To both extend and quantify the extent of this rDNA hypercondensation defect, we synchronized wildtype and *hsp82* deletion cells in G1 at 23°C, then released divided cultures into either 23°C or 37°C medium supplemented with nocodazole to arrest cells in preanaphase. Cell cycle progression and arrests were monitored using flow cytometry (Figure 4C). We then measured the axial rDNA loop length from three biological replicates. The results reveal that mitotic *hsp82* mutant cells shifted to 37°C contain significantly longer (roughly 30%) rDNA loops than wildtype cells shifted to 37°C (Figure 4D, E). Importantly, both wildtype and *hsp82* deletion cells exhibited similarly long loops at 23°C, further highlighting the unique role for Hsp82 in specifically driving hyperthermic-induced rDNA hypercondensation.

**Figure 4.**
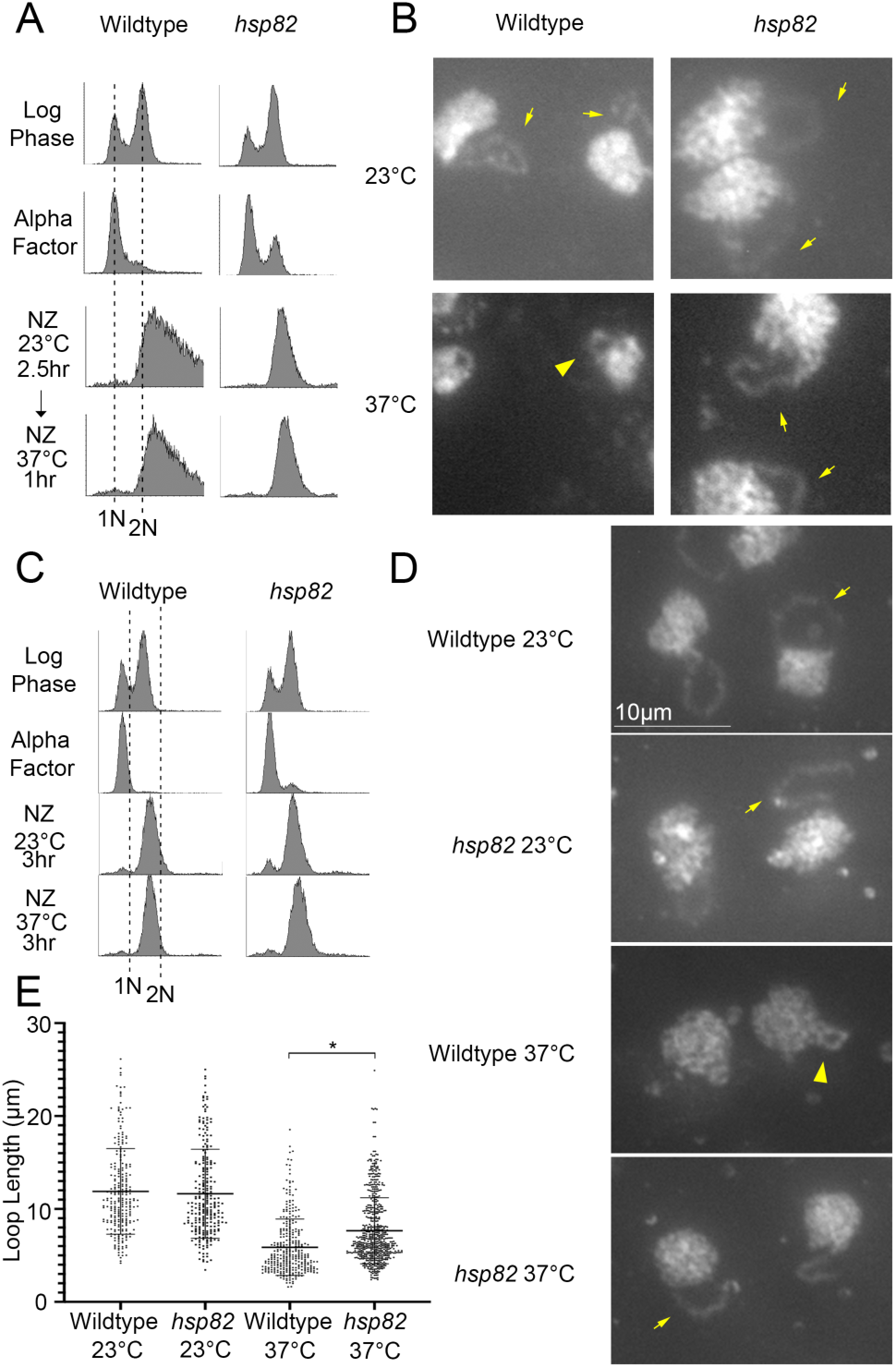
Hsp82 promotes hyperthermic-induced rDNA hypercondensation. A) Flow cytometer data documents DNA content for wildtype and *hsp82* synchronization (note that wildtype DNA profiles also appear in Figure 3A). Cells were maintained in nocodazole for 2.5 hours at 23°C, post-alpha factor synchronization/release, followed by an additional 1 hour incubation at 37°C. B) Chromosomal mass and rDNA loop structures, detected using DAPI, from cells obtained following a 1 hour shift up to 37°C. Yellow arrows indicate long rDNA loops. Yellow arrowheads indicate short rDNA loops. C) Flow cytometer data of DNA content for wildtype and *hsp82* synchronization. Cells were maintained in nocodazole for 3 hours at 23°C or 37°C, post-alpha factor synchronization/release. D) Chromosomal mass and rDNA loop structures, detected using DAPI, from cells obtained following a 3 hour incubation at 37°C. Yellow arrows indicate long rDNA loops. Yellow arrowheads indicate short rDNA loops. E) Quantification of the loop lengths of condensed rDNA in wildtype (YPH499) and *hsp82* null mutant (YDS203) cells. Data obtained from 3 biological replicates in which n values are 251 for Wildtype cells at 23°C, 254 for *hsp82* cells at 23°C, 323 for Wildtype cells at 37°C, 340 for *hsp82* cells at 37°C. Error bars represent standard deviation of each sample. Statistical analyses were performed using Tukey HSD one way ANOVA. P-Value = 0.001 indicates significant differences between the average loop lengths of wildtype cells (5.88 μm) versus the *hsp82* mutant cells (7.65 μm) at 37°C. Statistical significant differences (*) are based on P < 0.05. All micrographs are shown at the same magnification (see scale bar in D).

In yeast, Hsp90 family members include paralogs Hsp82 and Hsc82 that are 97% identical at the amino acid level, but exhibit differences in their expression (Kravats et al., 2018). Thus, it became important to determine the extent to which *hsc82* null cells phenocopy the hyperthermic-induced rDNA hypercondensation defect observed in *hsp82* null cells. Wildtype and *hsc82* deletion cells were synchronized in G1 at 23°C using alpha factor, then released into 37°C fresh medium supplemented with nocodazole for 3 hours to arrest cells in preanaphase as described above. Cell cycle progression and arrests were monitored using flow cytometry (Figure 5A). The results reveal that *hsc82* null cells exhibit a 40% increase in rDNA loop length, compared to wildtype cells (Figure 5B, C). A statistically significant increase (30%) in rDNA loop lengths was also observed in a second iteration in which we compared *hsc82* null cell rDNA loops lengths to those obtained from a BY4741 background strain (Supp. Figure 3). In combination, our combined findings from both *hsc82* and *hsp82* null cells, compared to 8 other heat-response proteins, reveal that Hsp90 function is required for thermic-induced rDNA hypercondensation.

**Figure 5.**
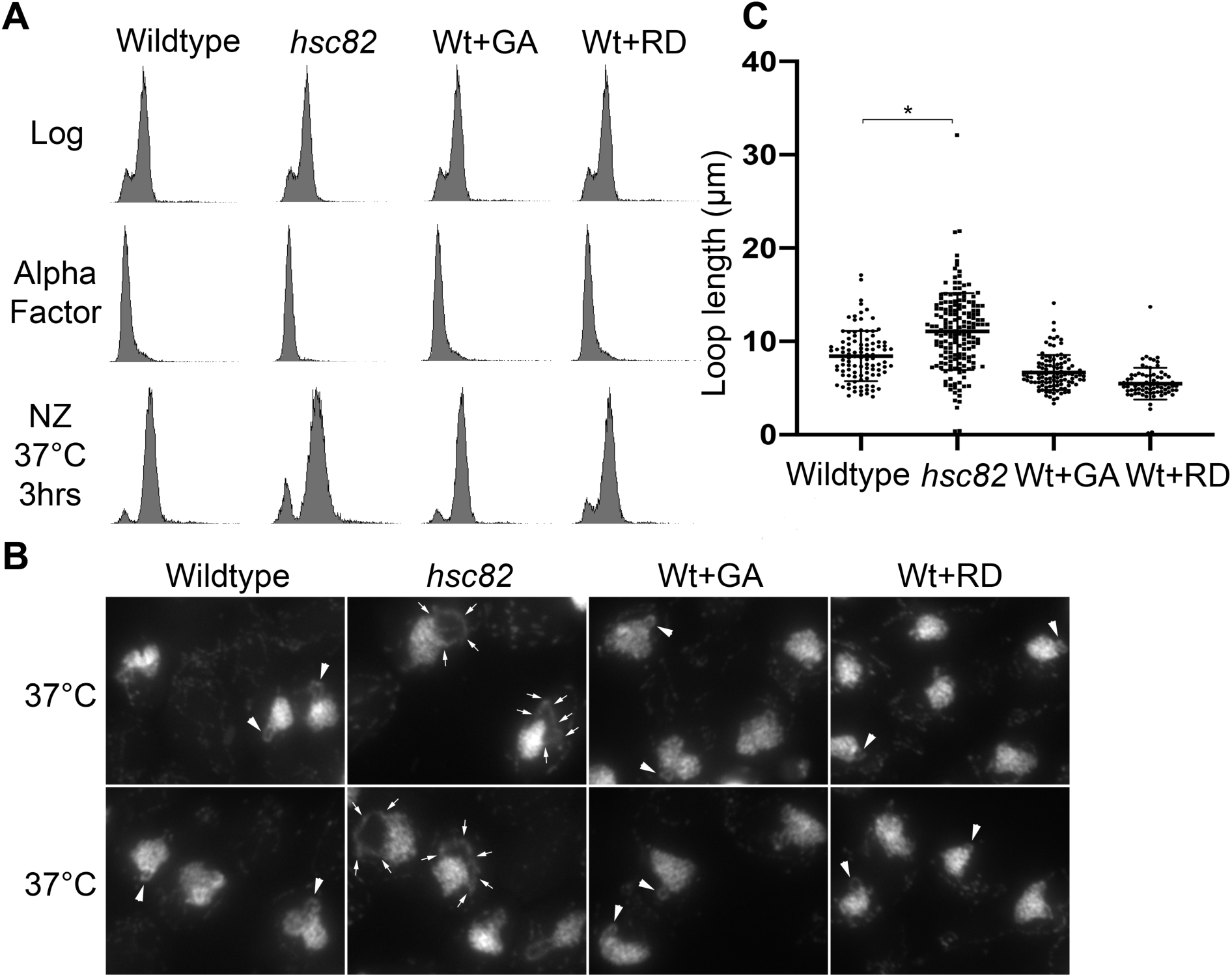
Impact of Hsc82 deletion and Hsp90 inhibitors on hyperthermic-induced rDNA hypercondensation. A) Flow cytometer data documents DNA content for wildtype (YPH499) and *hsc82* null cells (and see Supplemental Figure 3). For each, log phase cells were synchronized in G1 using alpha factor, then released into 37°C fresh medium supplemented with nocodazole (NZ) for 3 hours to arrest cells in preanaphase. Note that the culture of G1 synchronized wildtype cells was split into three aliquots (NZ+/- Geldanamycin (GA) or Radicicol (RD)), which is reflected by duplication of the Log and G1 DNA profiles for those treatments. B) Chromosomal masses and rDNA loop structures were detected using DAPI. White arrows indicate the track of rDNA long loops, arrowheads indicate rDNA short loops. C) Quantification of loop lengths of condensed rDNA in wildtype (YPH499) and *hsc82* null cells (YDS210). The quantified results are based on n values of 102 for Wildtype cells at 37°C and 170 for *hsc82* cells at 37°C. Error bars represent standard deviation of each sample. Statistical analyses were performed using Tukey HSD one way ANOVA. P-Value = 0.001 indicates significant differences between the average loop lengths of wildtype cells (4.21 μm) versus the *hsc82* mutant cells (5.53 μm) at 37°C. Statistical significant differences (*) are based on P < 0.05.

Hsp90 family members exhibit receptor/kinase signal transduction activities and are well-established ATP-dependent foldases that promote protein maturation and thermic tolerance by ensuring proper folding of client proteins (Khurana et al., 2018; Morán Luengo et al., 2019; Genest et al., 2019). In addition, however, Hsp90 family members also exhibit ‘holdase’ functions independent of ATP binding/hydrolysis that include structural or scaffolding roles (Genest et al., 2019; Hoter et al., 2018; Csermely et al., 1998). In this light, we were particularly intrigued by early EM studies through which Hsp90 was localized to the nucleolus and onto chromatin fibrils (Biggiogera et al., 1996; Ohtani et al., 1995). We thus decided to test whether inhibition of Hsp90 ATPase activity, while retaining Hsp90 scaffolding function, would adversely impact hyperthermic-induced rDNA hypercondensation. Geldanamycin (GA) and Radicicol (RD) are both potent Hsp90 inhibitors that bind the ATP binding pocket and preclude foldase activity (Chen et al., 2012; Wider et al., 2009; Theodoraki et al., 2012; Millson et al., 2014; Roe et al., 1999). Wildtype cells were synchronized in G1, using alpha factor, and then released into 37°C rich medium supplemented with nocodazole alone or further supplemented with either GA (40µM) or RD (20µM versus 40 µM for two experimental iterations) for 3 hours to arrest cells in preanaphase in the absence of Hsp90 ATPase activity. Cell cycle progression and arrests in this shift-up experiment were confirmed using flow cytometry (Figures 5A and Supp. Figure 3A). We then measured the axial rDNA loop length from two biological replicates in which each contains at least 100 cells. Neither Hsp90 inhibitor (GA or RD) had any adverse effect on rDNA hypercondensation (Figures 5B, C and Supp. Figure 3B, C). GA and RD entry into yeast and inhibition of Hsp90 are likely immediate with overt responses obtainable within minutes (Theodoraki et al., 2012; Tahbaz et al., 2001). We validated GA-dependent Hsp90 inhibition on the Hsp90 client protein Chl1, a DNA helicase that promotes Scc2/cohesin deposition onto DNA (Supp. Figure 4; Skibbens, 2004; Khurana et al., 2018; Tahbaz et al., 2001). In combination, these results provide intriguing evidence that Hsp90 proteins may play ATP-independent ‘holding’ activities that are critical for rDNA responses to thermic stress.

### Hmo1 negatively regulates mitotic rDNA condensation

During our screen of 10 HSP/C null cells function in hyperthermic-induced rDNA hypercondensation, we observed that *hmo1* null cells contained two distinct condensed rDNA loops – often appearing as rabbit ears (Supp. Figure 5A). The fact that isolates from this strain had diploidized was confirmed by flow cytometry (Supp. Figure 5B). Intriguingly, these *hmo1* null cells often failed to arrest with a 2N DNA content in response to medium supplemented with nocodazole (Supp. Figure 5B), suggesting that there are additional mutations that reside in the *hmo1* deletion strain (Winzeler et al., 1999; Giaever and Nislow, 2014; VanderSluis et al., 2014). Regardless of the aforementioned phenotypes, we noted that *hmo1* isolates also contained aberrant rDNA loop lengths (see below).

To further investigate Hmo1 function in hyperthermic-induced rDNA hypercondensation, we generated new *hmo1* null strains in the S288C wildtype background, confirming specific gene replacement by PCR (see Materials and Methods). We then synchronized wildtype and three independent *hmo1* null isolates for 3 hours (at 23°C or 37°C) in medium supplemented with nocodazole and then measured rDNA loop lengths for each of the preanaphase-arrested strain. Flow cytometry results document that each of our *hmo1* deletion isolates is haploid and arrests in response to nocodazole (Figure 6A). As expected, wildtype cells contained extended rDNA loops at 23°C and hypercondensed rDNA loops at 37°C (Figure 6B). Surprisingly, *hmo1* mutant cells contained significantly shorter loops at 23°C, compared to wildtype. Upon exposure to 37°C, however, the rDNA loops in *hmo1* null cells hypercondensed to a length similar to that exhibited by wildtype cells (Figure 6B, C). Thus, while mitotic *hmo1* cells contain increased levels of rDNA condensation at 23°C, a shift to 37°C does not promote rDNA hypercondensation beyond that observed in wildtype cells. Given that *hmo1* null cells exhibit elevated rDNA condensation in the absence of hyperthermic stress, we term Hmo1 a novel negative regulator of mitotic rDNA condensation.

**Figure 6.**
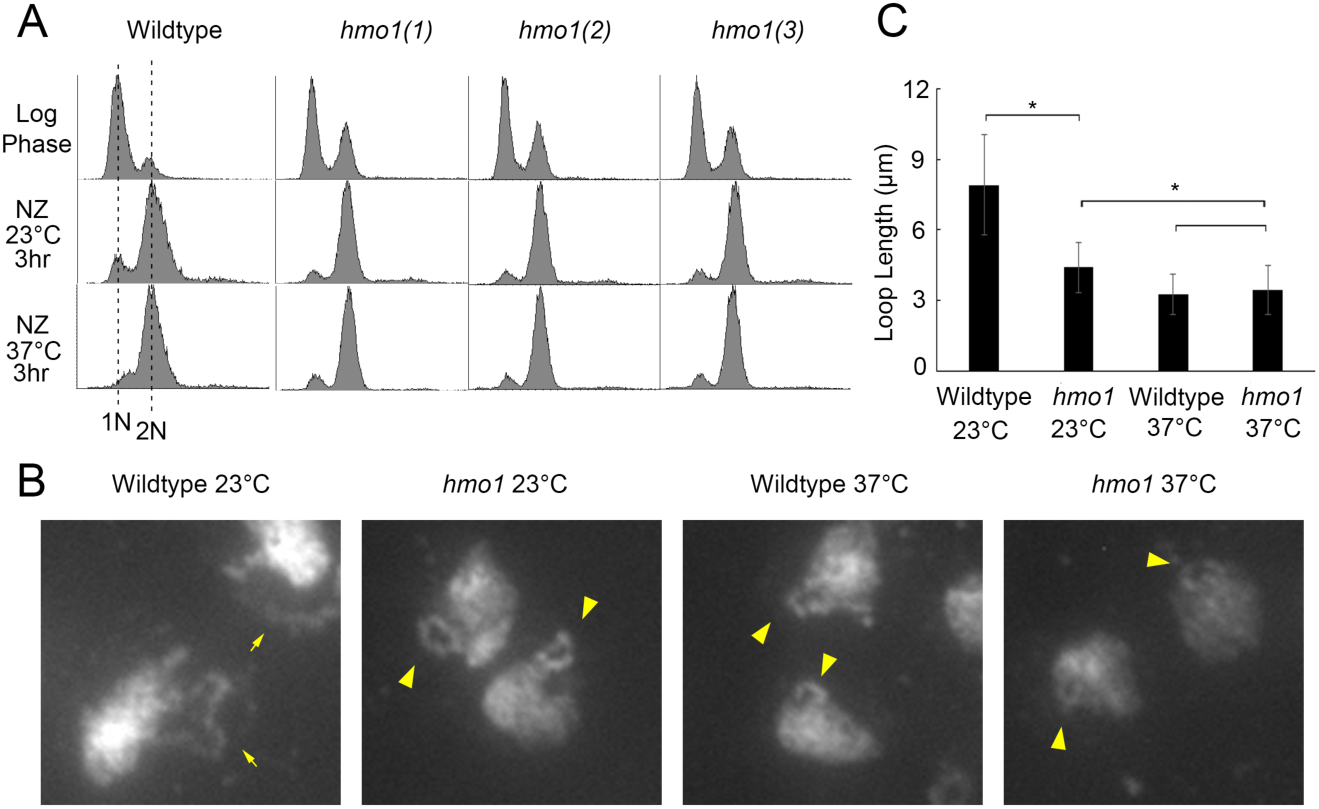
Hmo1 negatively regulates mitotic rDNA condensation. A) Flow cytometer data documents DNA content throughout the experiment. Log phase cultures were split and equal portions incubated for 3 hours at either 23°C or 37°C in fresh media supplemented with nocodazole. B) Chromosomal mass and rDNA loop structures detected using DAPI. Yellow arrows indicate long rDNA loops. Yellow arrowheads indicate short rDNA loops. C) Quantification of the loop lengths of condensed rDNA in wildtype (YPH499) and three *hmo1* deletion mutant (YBS3047, YBS3048, YBS3049) cells. Graphed values are based on n values of 100 for Wildtype cells at 23°C, 182 for *hmo1* cells at 23°C, 48 for Wildtype cells at 37°C, and 159 for *hmo1* cells at 37°C. Error bars represent standard deviation of each sample. Statistical analysis performed using Tukey HSD one way ANOVA. P-Value = 0.001 indicates there is significant differences between the average loop lengths of wildtype cells versus the *hmo1* mutant cells arrested at 23°C. P-Value = 0.001 indicates there is significant differences between the average loop lengths of *hmo1* mutant cells arrested at 23°C versus 37°C. P-Value = 0.756 indicates there is no significant differences between the average loop lengths of wildtype cells (3.24 μm) versus the *hmo1* mutant cells (3.44 μm) arrested at 37°C. Statistical significant differences (*) are based on P < 0.05.

## DISCUSSION

The nucleolus and rDNA are exquisitely tuned to both the cell cycle and external cues. For instance, rDNA prematurely condenses during interphase in response to starvation and also condenses in a stereotypic fashion during each entry of the cell into mitosis (Guacci et al., 2004; Li et al., 2006; Tsang et al., 2007). All of these structural changes require condensins with an additional role played by cohesins during mitotic condensation (Tsang et al., 2007; Wang et al., 2016; Guacci et al., 2004; Lopez-Serra et al., 2013; Skibbens et al., 1999; Tóth et al., 1999; Kueng et al., 2006; Hirano, 2012). Recently, we and others identified a novel form of hyper-condensation that occurs during mitosis in response to heat stress and thus far appears specific to the rDNA (Shen and Skibbens, 2017a; Matos-Perdomo and Machín, 2018a). The first major finding of the current study is that rDNA hyperthermic-induced hypercondensation occurs independent of condensin and cohesin recruitment/dissociation. This surprising result suggests that the last several decades of research into rDNA structure analyses remain narrowly focused on SMC complexes and that our understanding of chromatin structure regulation remains incomplete.

Recently, a condensation end-state (also referred to as hypercondensation) was reported to occur during anaphase and that required additional condensin recruitment (de los Santos-Vas et al., 2017). A second major finding of the current study, based on analyses of anaphase promoting complex mutant strains, is that this normal chromatin condensation end-state reported by de los Santos-Vas and colleagues is distinct from the thermic-induced rDNA hypercondensation that occurs during preanaphase. Moreover, new findings reported here document that the significant rDNA axial loop shortening that occurs during preanaphase proceeds in the absence of additional condensin recruitment. In support of these distinct mechanisms, we note that thermic-induced rDNA hypercondensation is rapidly reversible, while the proteolytic mechanism that underlies anaphase onset is not (Amon et al., 1994; Cohen-Fix et al., 1996; Ulhmann et al., 1999; Shen and Skibbens, 2017a).

A third major finding of the current study is that mutation in either *HSP82* and *HSC82*, which encode members of the Hsp90 HSP/C family, results in defective hyperthermic-induced rDNA hypercondensation. Here, we consider several possibilities regarding the mechanism through which HSP/C impact thermic-induced rDNA hypercondensation. Given the well-established role for HSP/C in stabilizing or refolding client proteins (Khurana et al., 2018; Morán Luengo et al., 2019; Genest et al., 2019), one plausible mode of action is through stabilization of an as yet undefined thermic-sensitive client protein required for rDNA hypercondensation. This client is unlikely to include cohesin or condensin, given that thermic stresses in HSP/C mutant cells result in increased rDNA axial loop lengths but not loss of either loop morphology (loops transitioning to puffs) or sister chromatid cohesion (one loop transitioning to two loops) (Shen and Skibbens, 2017a; Matos-Perdomo and Machín, 2018a). In contrast, cohesin inactivation quickly results in rDNA puff structures and cohesion loss (Guacci et al., 1994; Guacci et al., 1997; D’Ambrosio et al., 2008; Lopez-Serra et al., 2013; Tong and Skibbens, 2015; Lavoie et al., 2002; Lavoie et al., 2004; Shen and Skibbens, 2017a). We further note that thermic-induced hypercondensation appears to effect rDNA specifically (Shen and Skibbens, 2017a), while cohesins and condensin impact chromatin architecture genome-wide (Shen and Skibbens, 2017b; Rudra and Skibbens, 2012; Skibbens, 2004; Mayer et al., 2004; Xu et al., 2007; Borges et al., 2013; Samora et al., 2016). Importantly, potent Hsp90 ATP-binding/hydrolysis inhibitors Geldanamycin and Radicicol (Chen et al., 2012; Wider et al., 2009; Theodoraki et al., 2012; Millson et al., 2014; Roe et al., 1999) both failed to adversely impact hyperthermic-induced rDNA (current study). Thus, a novel and exciting possibility is that Hsp90 family members play a direct structural role in hypercondensing rDNA in response to heat stress. This model is supported both by EM studies that localize Hsp90 to the nucleolus and also by circular dichroism spectra studies that Hsp90 induces *in vitro* a more condensed chromatin state in rat liver cells (Csermely et al., 1994; Ohtani et al., 1995; Biggiogera et al., 1996). A third possibility is that Hsp90 ‘holdases’ regulate factors that in turn promote rDNA hypercondensation. For instance, Hsp82 exhibits synthetic growth defects with histone (H2B), histone variant (H2A.Z), histone modifiers and chromatin remodeling complexes (Dep1, Eaf1,7, Gcn5, Gis1, Hda2,3, Pho23, Rco1, Rtt109, Sap30, Set2 and Swi3) (Millson et al., 2005; Zhao et al., 2005; McClellan et al., 2007). Such histone modification cascades (including deacetylation, phosphorylation and tail-tail interaction of adjacent histones) may promote rDNA hypercondenses during mitosis (Wilkins et al., 2014). Future efforts are required to resolve the issue of whether Hsp82 and Hsc82 directly impose rDNA structure or promote (through foldase or holdase activities) other factors to induce hyperthermic-induced rDNA hypercondensation.

While the results presented here argue against a role for either cohesin or condensin deposition/release in driving rDNA hypercondensation, we cannot rule out a model in which post-translational modifications alter the looping activities of these SMC complexes. For instance, it is well established that condensin phosphorylation promotes mitotic condensation (Kakui and Uhlmann, 2018; Kinoshita and Hirano, 2017; Kalitsis et al., 2017). Cdc28 is the cyclin-dependent kinase that phosphorylates condensin and triggers condensation during prophase (Hadwiger et al., 1989; Surana et al., 1991). It is thus notable that Hsp82 and Cdc28 physically interact (Zarzov et al., 1997), providing a mechanism through which Hsp82-recuitment of CDK might activate condensins without altering deposition/release dynamics. Intriguingly, Cdc14 phosphatase also impacts rDNA condensation and physically interact with Hsp82, potentially revealing a complex interplay between post-translational modification of condensins and rDNA architecture prior to anaphase (de los Santos-Vas et al., 2017; Sullivan et al., 2004; Woodford et al., 2016). Cohesin modification, in the absence of deposition/dissociation, might also contribute to rDNA condensation upon thermic stress during preanaphase. For instance, *hsp82* mutants exhibit synthetic growth defects in combination with mutation of the cohesin acetyltransferase *ECO1* (Skibbens et al., 1999; Tóth et al., 1999; Millson et al., 2005; Borges et al., 2013; Rolef Ben-Shahar et al., 2008). While there is a paucity of evidence that physically links Eco1 to Hsp82, it remains formally possible that Hsp82 promotes rDNA hypercondensation through Eco1 acetylation of cohesin subunits. In contrast to reports that Hsp82 promotes sister chromatid cohesion through stabilization of Chl1 DNA helicase (Khurana et al., 2018), which in turn promotes both Scc2 and cohesin binding to DNA (Skibbens, 2004; Rudra and Skibbens, 2012; Shen and Skibbens, 2017b; Mayer et al., 2004; Xu et al., 2007; Borges et al., 2013; Samora et al., 2016), we find no evidence in the current study that Chl1 (or changes in cohesin/condensin dynamics) adversely impacts rDNA hypercondensation. Future studies will be required to ascertain the extent to which, histone, cohesin and/or condensin modifications promote hyperthermic-induced rDNA hypercondensation in an Hsp90-dependent manner (Figure 7).

**Figure 7.**
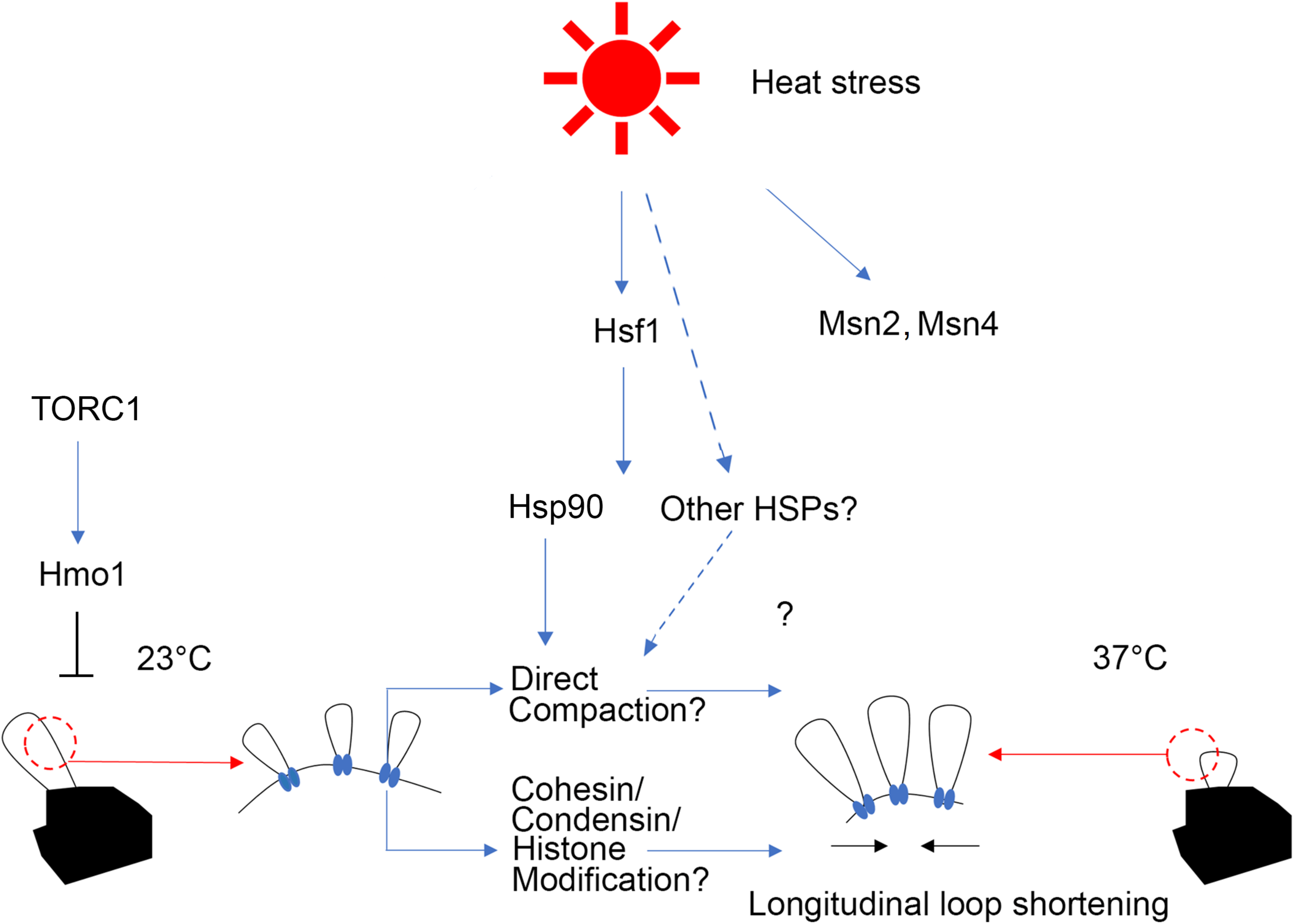
Possible mechanisms of Hsp90-dependent hyperthermic-induced rDNA hypercondensation. Hsp90 functions in hyperthermic-induced rDNA hypercondensation possibly through 1) direct interaction of rDNA, 2) recruitment of enzymes that post-translationally modify cohesins, condensins or histones or 3) by stabilizing client proteins that perform novel roles in rDNA structure (not shown). Here, we suggest that Hmo1 may be involved in TORC1 signaling that antagonize rDNA condensation. Hmo1 inhibits hyperthermic-induced rDNA hypercondensation such that rDNA hypercondenses in the absence of Hmo1. The impact of rDNA axial hypercondensation in generating transcriptionally active lateral loops is hypothetical, but consistent with increased transcription at 37°C.

A fourth major finding of the current study is that the High Mobility Group protein Hmo1, that is involved in TOR signaling, negatively regulates mitotic rDNA condensation. Inhibition of TOR by rapamycin causes nucleolar contraction, condensin loading on to rDNA during interphase, and also promotes rDNA hypercondensation in both pre-anaphase and anaphase-arrested cells (Tsang et al., 2007; de los Santos-Vas et al., 2017; Matos-Perdomo and Machín, 2018a; Matos-Perdomo and Machín, 2018b). Thus, TOR inhibition might trigger increased condensation of rDNA through Hmo1. Despite the increased condensation state of rDNA that occurs at 23°C in *hmo1* deletion cells, rDNA remains competent to hypercondense upon a temperature shift to 37°C. This latter hypercondensed state is similar to that exhibited by wildtype cells at 37°C, suggesting that Hmo1 antagonizes rDNA compaction through a mechanism separate from Hsp90-dependent hyperthermic-induced rDNA hypercondensation. Conversely, Hmo1 promotes rDNA transcription and triggers DNA bridging and looping that contributes to higher-order architectures (Wang et al., 2016; Divakaran et al., 2014; Albert et al., 2013). Thus, on the one hand, Hmo1 promotes condensation (in a condensin-dependent fashion) during interphase in response to nutrient starvation while, on the other hand, antagonizes mitotic condensation (Wang et al., 2016 and the current study).

It is remarkable that a GO-term based bio-informatic screen, in which we limited analyses to only ten candidates, turned up two factors: Hsp82 that is critical for hyperthermic-induced rDNA hypercondensation and Hmo1 that negatively regulates canonical mitotic condensation. We were able to validate our results for the role of Hsp82 by exploiting the highly-conserved paralog Hsc82. While the mechanism through which HSP/C impacts thermic-induced rDNA hypercondensation remains unknown, this is the first report that HSP/C support this process and suggest that future endeavors will continue to uncover novel roles for HSP/C function in chromatin structure. Our results further document that single mutant *hsp82* and *hsc82* null cells are only partially inhibited for hyperthermic-induced rDNA hypercondensation. Synthetic lethality and/or severe growth defects precluded testing *hsp82 hsc82* double mutants (McClellan et al., 2007; Hainzl et al., 2009). Given that there are more than 1000 genes involved in the heat shock response in budding yeast (Morano et al., 2012), we anticipate that the identification of redundant pathways will uncover novel mechanisms through which HSP/C factors help promote alterations in chromatin structure.

## MATERIALS AND METHODS

### Yeast strains and strain construction

Saccharomyces cerevisiae strains used in this study are listed in Supplementary Table 1 (SI). Primers used to verify gene deletions within the Knockout collection are available upon request.

### rDNA condensation assay

A streamlined condensation assay is adapted from a published FISH protocol (Guacci et al.,1997; Shen and Skibbens, 2017a). Briefly, cells were arrested at preanaphase and fixed by paraformaldehyde for 2hr at 23 °C. Cells were washed with distilled water and resuspended in spheroplast buffer (1M sorbitol, 20mM KPO4, pH7.4), then spheroplasted by adding beta-mercaptoethanol and Zymolyase T100 and incubating for 1 hour at 23°C. Resulting cells were added to poly-L-lysine coat slides, treated with 0.5% Triton X-100, 0.5% SDS, and dehydrated in 3:1 methanol:acetic acid. Slides were stored at 4°C until complete dry, then cells were treated with RNase in 2XSSC buffer (0.3M NaCl, 30mM Sodium Citrate, pH7.0), dehydrated and denatured under 72°C following cold ethanol wash. DNA mass were detected by DAPI staining and assayed under microscope. Cell cycle progression were confirmed by detection of DNA content using flow cytometry as described (Tong and Skibbens, 2015).

### Chromatin Immunoprecipitation and ChIP primers

ChIP was performed as previously described (Rudra and Skibbens, 2012), with the following modifications. Cells were cultured to log phase with OD600 1.0 to 1.2, then incubated at 23 °C in rich YPD medium supplemented with alpha-factor for 2.5 hr. The resulting cells were collected, washed and then resuspended in fresh YPD supplemented with nocodazole, incubated at 23 °C or 37°C for 3 hours, and then fixed in 1% formaldehyde for 20 mins. Cells were then harvested, spheroplasted and lysed. Cells lysates were sonicated on ice for 6 cycles of 10 seconds. The suspension was centrifuged and diluted 1:10. The diluted suspension was then centrifuged and the supernatant was collected as the chromatin solution. Smc2 enrichment was obtained by incubating chromatin solution with EZ-View Red Anti-HA affinity matrix (Sigma) overnight at 4°C, the background control was obtained in a similar manner by incubating the same batch of chromatin solution (isogenic strain expressing Smc2-HA) with EZ-View Red Anti-Myc affinity matrix (Sigma, used as beads only control) overnight at 4°C. Beads were collected by centrifugation, washed and the remaining bead-bound proteins harvested using 1%SDS; 0.1 M NaHCO3. DNA-protein crosslinks were reversed in 5 M NaCl for 4 hr at 65°C. DNA precipitation from the resulting lysate was performed by overnight incubation at −20°C in 70% ethanol. Precipitates were extracted in series using 25:24:1 phenol:chloroform:isoamylalcohol and pure chloroform prior to reprecipitation of DNA overnight at −20°C in 70% ethanol. DNA was resuspended in TE buffer and analyzed by PCR using rDNA primers previously described (Johzuka and Horiuchi, 2009; Thattikota et al., 2018). PCR products were resolved using 1% agarose gels, and histograms of pixel densities quantified in Photoshop. Smc2 enrichment was calculated as the ratio of HA pull down (ChIP) minus Myc pull-down (background) all over total chromatin input.

### Statistical Analyses

Tukey HSD one way ANOVA test were used to assess the statistical significance (P < 0.05).

