## Supplementary figures and images for "Promotion of hyperthermic-induced rDNA hypercondensation in *Saccharomyces cerevisiae*"

### Supplemental Figure 1

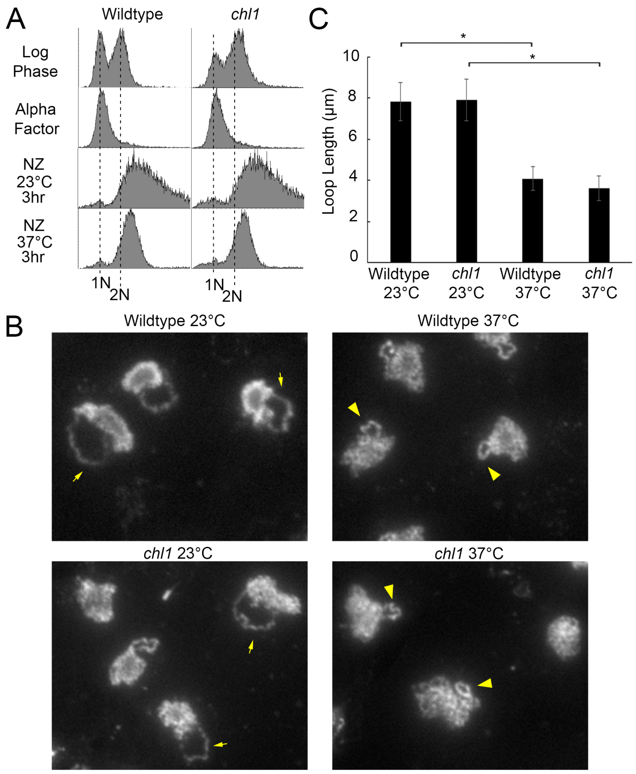

### Supplemental Figure 2

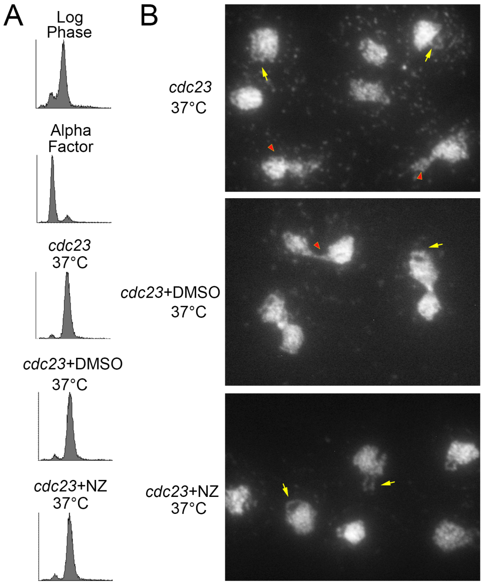

### Supplemental Figure 3

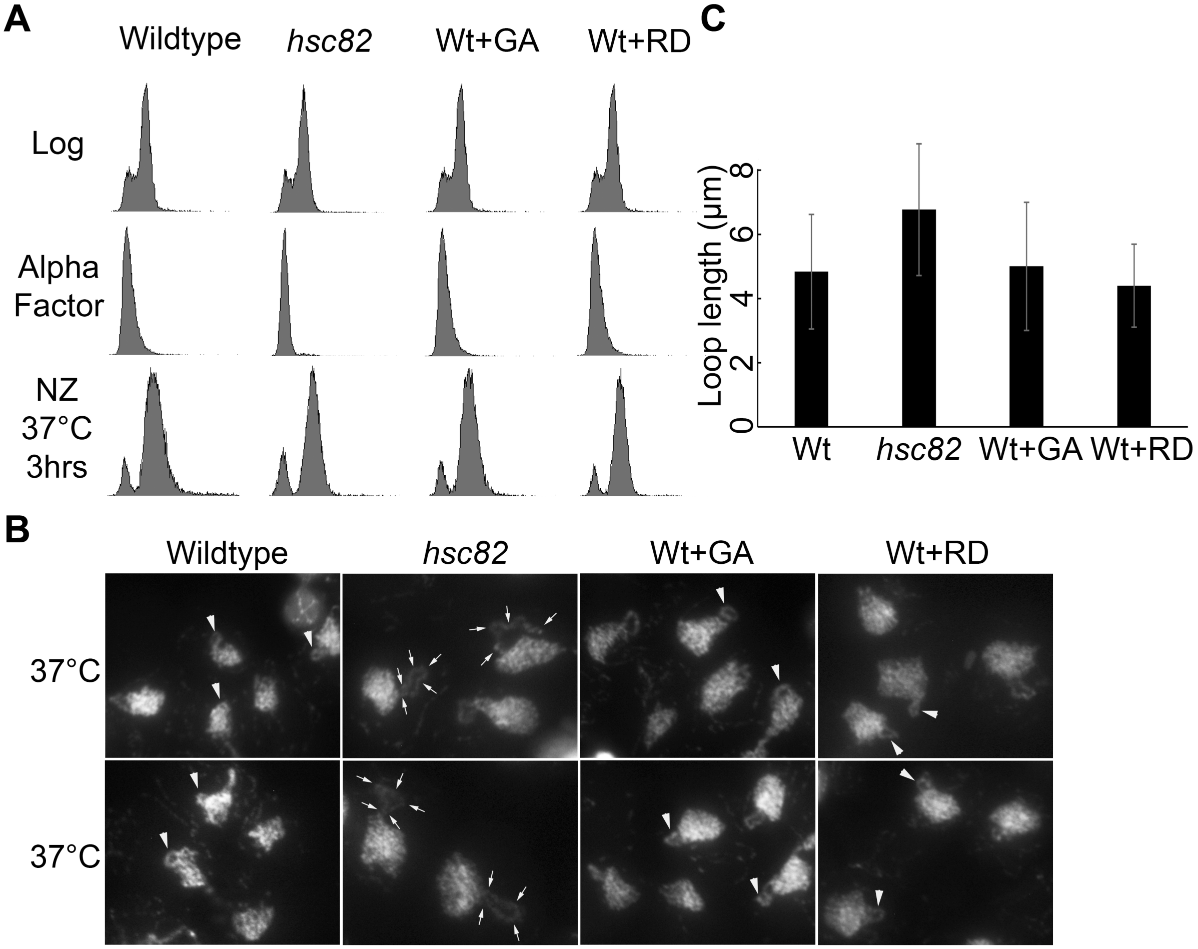

### Supplemental Figure 4

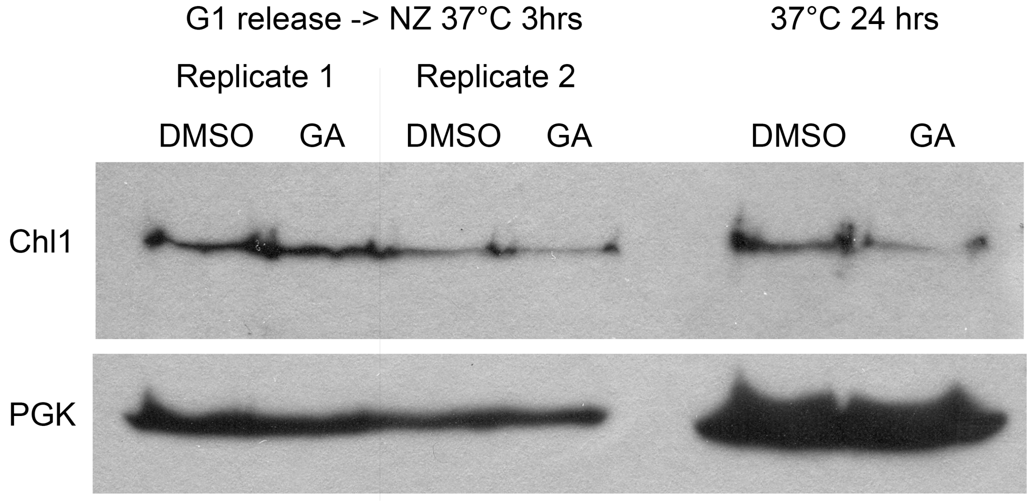

### Supplemental Figure 5

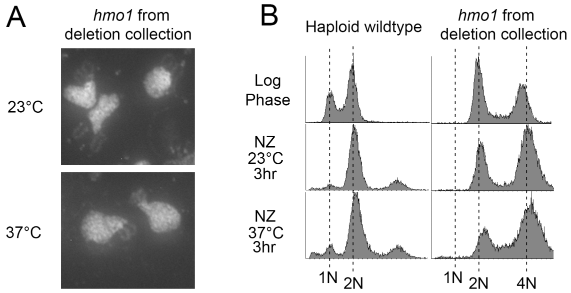
