## Supplemental Figure legends for "Promotion of hyperthermic-induced rDNA hypercondensation in *Saccharomyces cerevisiae*"

**Supplemental Figure 1. Chl1 DNA helicase does not function in hyperthermic-induced rDNA hypercondensation.** A) Flow cytometer data of DNA content. Log phase cultures were synchronized in G1 using alpha factor and aliquots released into either 23°C or 37°C fresh media supplemented with nocodazole and incubated for 3 hours. B) Chromosomal mass and rDNA loop structures detected using DAPI. Yellow arrows indicate long rDNA loops. Yellow arrowheads indicate short rDNA loops. All field of views are shown at equal magnification. C) Quantification of condensed rDNA loop lengths in wildtype (YPH499) and chl1 deletion mutant cells (YBS1141). Data shown was obtained from 3 biological replicates with at least 100 cells for each strain analyzed per replicate. Error bars represent standard deviation of each sample. Statistical analysis was performed using a Student's T-test. P-Value = 0.0053 indicates a significant difference between the average loop lengths of wildtype cells at 23°C versus 37°C. P-Value = 0.0030 indicates significant differences between the average loop lengths of chl1 mutant cells at 23°C versus 37°C. Statistically significant differences (*) are based on P < 0.05. Wildtype rDNA loop measurements adapted from Figure 1E, Shen and Skibbens 2017a, under the terms of the Creative Commons Attribution-NonCommercial-NoDerivatives License.

**Supplemental Figure 2. rDNA hypercondensation can be induced prior to anaphase onset.** A) Flow cytometer data of DNA content. Log phase cultures of cdc23-1 were synchronized in G1 using alpha factor and aliquots released into fresh medium along or medium supplemented with DMSO (+DMSO) or nocodazole (+NZ) and incublated for 3 hours at 37°C. B) Chromosomal mass and rDNA loop structures were detected using DAPI. All field of views are shown at equal magnification. Red arrows indicate distorted chromatin structure. Yellow arrows indicate hypercondensed rDNA loops. Hypercondensed rDNA loops are clearly present in cells that harbor temperature-sensitive alleles of cdc23-1 (FLY83/H23C1AX) at all three conditions.

**Supplemental Figure 3. Independent confirmation that Hsp90 protein**

**promotes hyperthermic-induced rDNA hypercondensation and the impact of**

**Hps90 inhibitors.** A) Flow cytometer data documents DNA content for wildtype

(BY4741) and hsc82 null cells. Log phase cells were synchronized in G1 using alpha

factor, then released into 37°C fresh medium supplemented with nocodazole for 3 hours

to arrest cells in preanaphase. Wildtype cells were further treated with Geldanamycin

(GA) or Radicicol (RD), in addition to NZ, which is reflected by duplicated Log and G1

DNA profiles for those treatments. B) Chromosomal mass and rDNA loop structures

detected using DAPI. White arrows indicate the track of rDNA long loops, arrowheads

indicate rDNA short loops. C) Quantification of loop lengths of condensed rDNA in

wildtype (BY4741) and hsc82 null cells (YDS210). At least 100 cells for each strain were

analyzed. Error bars represent standard deviation of each sample. Statistical analysis

performed using Tukey HSD one-way ANOVA. P-Value = 0.001 indicates significant

differences between the average loop lengths of wildtype cells versus the hsc82 mutant

cells at 37°C. Statistically significant differences (*) are based on P < 0.05.

**Supplemental Figure 4. Geldanamycin inhibits Hsp90 ATPase activity, resulting in Chl1 degradation.** Cells (YBS1129) expressing MYC-tagged Chl1 were exposed to short (3 hours) versus long (24 hours) incubations in 37°C fresh medium supplemented with nocodazole and either vehicle (DMSO) or 40µM GA, post G1 synchronization/release. As previously reported, cells exposed to 37°C in medium supplemented with nocodazole (NZ) and Geldanamycin (GA) for short durations (3 hrs) retain relatively high levels of Chl1. In contrast, cells exposed to GA for an extended period of time (24 hrs), contain dramatically reduced levels of Chl1, compared to DMSO vehicle. Phosphoglycerate kinase (PGK) levels are shown as an internal loading control.

**Supplemental Figure 5. Diploidization of hmo1 strain obtained from the yeast deletion collection.** A) Chromosomal mass and rDNA loop structures detected using DAPI. The majority of DNA masses exhibit two rDNA loops. B) Flow cytometer of DNA content. Log phase wildtype (YDS202) and hmo1 null strains (obtained from yeast deletion collection – a generous gift from Prof Greg Lang) were split and equal portions placed in fresh medium supplemented with nocodazole and incubated for 3 hours at either 23°C or 37°C.
